# Plant age and genotype explain variation in the microbiome of natural *Lotus corniculatus* populations

**DOI:** 10.1101/2025.05.02.651871

**Authors:** Frank Reis, Katrina Lutap, Jun Hee Jung, Anna Roeder, Christiane Roscher, Walter Durka, Eric Kemen, Oliver Bossdorf

## Abstract

In natural populations, plants are associated with a huge diversity of bacteria, fungi and other microbes. There is usually substantial microbiome variation between different plant individuals and populations, and the drivers of this variation are still poorly understood, particularly in wild plants. Here, we were interested in the potential of plant genotype and plant age to explain intraspecific variation in the plant microbiome of *Lotus corniculatus*. In seven natural populations, we genotyped a total of 168 individuals over four years, determined their ages through growth ring analysis, and then sequenced their root, shoot, flower and seed microbiomes. We found that plant genotypes differed both in the diversity and composition of microbes, and that some microbial taxa were associated with particular plant genotypes. The genotype effects tended to be strongest and most consistent for plant-associated bacteria, with the largest plant genotype differences in the microbiome diversity of flowers and seeds. We found less evidence for an effect of plant age on microbiome diversity: the age of plants explained variation in fungi diversity, and it was associated with the abundance of several microbial taxa. Our study indicates that the genotype of a plant and - to a lesser degree its age - influences the diversity and composition of plant-associated microbiota, even in complex natural environments.

**Importance:** The plant microbiome plays a key role in important plant functions such as pathogen resistance, nutrient uptake, and stress tolerance. To fully understand these processes, it is essential to identify the factors that drive microbiome variation. Most research to date has focused on model or crop species under controlled conditions, leaving open questions about the drivers of microbiome diversity and composition in natural populations. In this multi-year field study, we examined how plant genotype and age shape the microbiomes of *Lotus corniculatus*. We show that the plant genotype strongly influences microbial diversity and composition, while plant age has subtler but still important effects, particularly on fungal communities. To our knowledge, this is the first study to link plant age with microbiome variation in natural populations. These findings demonstrate that plant traits can shape the microbiome even in complex natural environments.

## Introduction

Plants are colonized by a wide range of bacterial, fungal and eukaryotic microorganisms collectively called the plant microbiome. The diversity and composition of this microbiome is strongly shaped by the plant species (1, 2) and the surrounding microbial community, especially the soil microbiome (3, 4), but there are many other biotic and abiotic factors that are known to influence plant microbiome variation among and within species, including climate (5, 6), soil characteristics (7), land use (8, 9), herbivory (10, 11) or plant diseases (12–16). Besides this plethora of external factors, the plant microbiome can also be influenced by the plant itself. Variations in gene expression and plant traits can result in changing physical and chemical properties of the plants, and thus habitat conditions for the microbes, which in turn can alter microbiome diversity and composition. Previous studies have shown that the plant microbiome indeed varies across different plant tissue types such as roots, shoots or flowers (17–19) and also between the inside (endophytic) and outside (epiphytic) microbiome (19, 20). Two other potentially important factors creating microbiome variation among plants of the same species are their genetic differences, and their differences in age or developmental stage.

It is well established that the genetic variation in plants can change the pathogen resistance (21–23), and there is also good evidence that it can influence the overall composition of the plant microbiome, e.g. in soybean (24), potato (10), black cottonwood (25). While these studies were all conducted under controlled lab conditions, there is also evidence from more natural common garden experiments, e.g. for plant genotype effects on the microbiome of *Medicago truncatula* (26) and *Boechera stricta* (27). So far the research on these questions has been largely restricted to crops and model species, and to - usually short-term - experiments, whereas studies on wild plants and natural populations remain rare. Still, extrapolating lab results about plant-microbe interactions to natural environments is often challenging (28) because in natural populations plant microbiomes are influenced by a much larger complexity of biotic and abiotic factors. A true understanding of plant genotype effects, and their relative importance, therefore requires to also study plant genotype-microbiome relationships in natural populations.

Another potentially important but so far very little studied intrinsic factor creating intraspecific variation in plant microbiomes is the age or developmental stage of a plant. As a plant grows and develops, it undergoes physiological and biochemical changes that influence important functional traits, such as defense against herbivory (29, 30) or immune responses to pathogens (31). These changes across different life stages will inevitably also influence plant-microbiome interactions (32). For example, the root-associated microbiome of rice varies between the vegetative phase and later life stages (33), and very similar results were found for soybean (34) and sugarcane (35). For annual plants like rice or soybean such a comparison between vegetative and reproductive phase covers their entire life cycle. However, many plants, in particular wild ones, are perennial, which creates additional possibilities for age-related microbiome variation e.g. through slower successional or competitive replacement processes. So far, there are hardly any studies of perennial plants examining relationships between plant age and plant microbiome. An exception is the study by Wagner et. al (2016) who found substantial changes in root (and to a lesser extent leaf) bacterial communities when comparing two and four year old *Boechera stricta* plants in a common garden, suggesting that significant microbiome changes can also occur at later life stages of perennials. Considering that most plants are perennial, we clearly need more studies relating plant age to plant microbiome in longer-lived plants.

Understanding the influences of plant age and genotype on microbiome composition may be important not only from a fundamental scientific perspective but also for agriculture and plant conservation, for instance if it helps to better manage beneficial crop microbiomes that increase yield production, stress tolerance or pathogen resistance. The last point is particularly important given that pathogens are a huge threat to global food security (36). More generally, understanding the drivers and consequences of plant microbiome diversity may support the management of plant populations in rapidly changing environments, and the effects of plant age and genotype on plant microbiomes - particularly under realistic conditions - are important elements of this.

Here, we combined amplicon-based metagenomics with growth ring analyses for plant age determination and genotype, and the diversity and composition of plant microbiomes within and among natural plant populations. Our study organism was *Lotus corniculatus*, a widely distributed perennial legume that grows naturally in a broad range of environments in temperate Eurasia and is an important food source for many pollinators, such as bumblebees, honeybees, and many wild bees as well as butterflies, flies and beetles (37–39). As a nitrogen fixer it is part of a complex plant-microbe symbiosis influencing the nutrient dynamics of grassland ecosystems (40), and it therefore plays an important role in agriculture. As *L. corniculatus* is known to have a broad life span range at least up to 14 years (41, 42), and it is known to harbor significant intraspecific genetic diversity (43, 44), the species is a suitable system for asking questions about effects of plant age and genotype. Specifically, we wanted to answer the following questions:

1. Do plant age and genotype affect the diversity of the plant microbiome, and if yes how do these effects vary between different plant organs and microbial groups?
2. Are plant age and genotype related to the composition (beta diversity) of the plant microbiome, and how are these relationships influenced by the plant organs and microbial groups?
3. Which specific microbial taxa are most differentiated between plants of different ages and genotypes?

To answer these questions, we collected *L. corniculatus* plants from seven semi-natural grasslands in the Swabian Jura region in Germany over four consecutive years, and we then determined the ages and genotype of all plants, and analyzed their bacterial, fungal and eukaryotic microbes separately for roots, shoots, flowers and seeds.

## Methods

### Plant material

We collected *Lotus corniculatus* plants from seven extensively managed meadows in the Swabian Alb region of southern Germany (Fig. 1a) for four consecutive years from 2018 to 2021. Sampling was done in the late summer (August/September), when the plants were still flowering but already carried some ripe fruits, so that both flowers and seeds could be sampled. In each of the seven populations, we carefully excavated six entire plants, resulting in a total of 4 years x 7 populations x 6 plants = 168 plants. In the lab, we separated the plants into roots, shoots, flowers and seeds. To surface sterilize all plant parts we washed them by shaking in Falcon tubes (50 ml for roots and shoots; 25 ml for flowers) and 1.5 ml Eppendorf tubes for seeds for 30 s with autoclaved ultrapure water (UPW), for 60 s with epiphyte wash solution (0.1 % Triton X-100 in 1x TE buffer), for 30 s with 80% EtOH and for 30 s with 2 % NaOCl. After that, we rinsed the samples three times (seeds: five times) for 10 s with UPW, and froze them.

**Figure 1.**
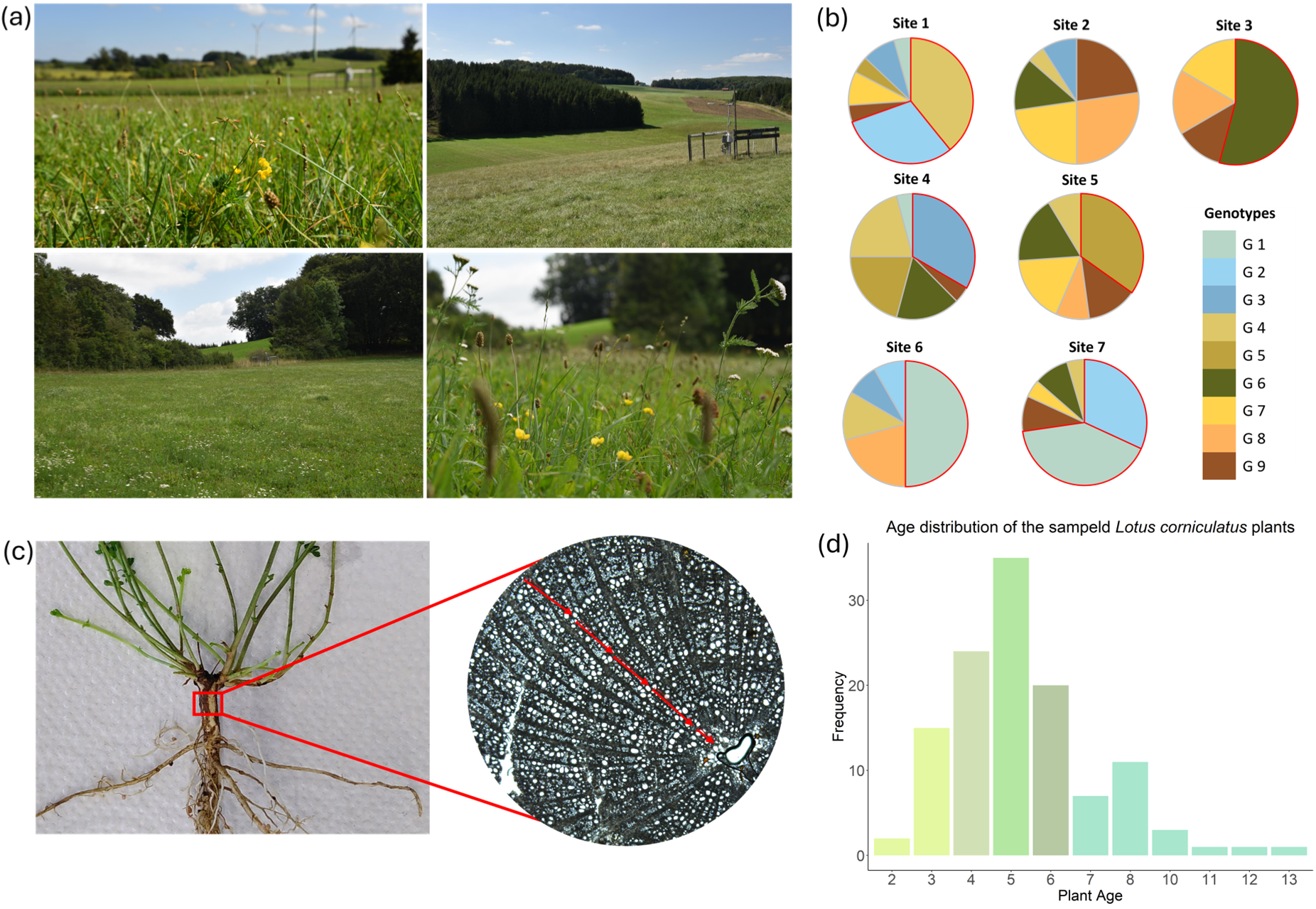
Sampling region, genotypic diversity, and determination of plant age. **a)** Photos of four of the sampling sites in southwest Germany. **b)** Frequencies of *Lotus* genotypes in the seven studied populations. The red borders indicate cases where genotypes are significantly overrepresented in specific populations. **c)** The root crown of a *Lotus* plant, and its cross-section, with the growth rings used to assess plant age. **d)** Age distribution of all sampled plants based on herb chronology. The colors indicate the five age groups we created for the statistical analysis.

### Determination of plant ages

To determine the age of the sampled plants, we cut an approx. 3 cm long piece of the oldest part of the plant, i.e. the root collar as the transition zone between the primary root and the germination stem, from all plants and preserved them in 70% ethanol. In the lab, we placed these samples into a microtome (Reichert, Vienna, Austria) and sliced them into transverse sections. Very small samples with a diameter smaller than 3 mm were clamped using a piece of wood. To harden the soft tissue and keep the samples moist for cutting, the sample and the microtome blade were brushed with 99% ethanol during the cutting process. The microtome sections were then fixed on a microscope slide in a 1:1 mixture of water and glycerol and covered with a cover glass. We used a microscope to count the number of growth rings in the xylem, with ring boundaries defined by marginal parenchyma, semi-ring porosity or both (45) (Fig. 1b). Where necessary we used polarized light to improve the visibility of the growth rings. Since the growth rings were not equally distinct in all specimens, we classified the distinctness of the rings into 1 = very good, 2 = intermediate, 3 = poor and 4 = no annual rings visible (Tab. S1) and used only classes 1 to 3 for our analyses. For the statistical analysis we decided to group the plants into five age groups since the low number of replicates for the very young (2–3 years) and very old plants (6–13 years). Therefore we grouped the plants into age categories: younger than 4, 4, 5, 6, and older than 6 years (Fig. 1d).

### Plant genotyping

To genotype the sampled plants, and assess their relatedness, we employed ddRAD (46), a reduced-representation sequencing approach. We extracted plant DNA from all shoot samples using the QIAGEN DNeasy Plant Mini Kit (QIAGEN, 2019). We then double-digested the DNA and followed the library preparation protocol of Peterson et al. (2012) (46) with small modifications as given in Durka et al. (2025) (44). Multiplexed samples were sequenced paired-end (PE 150) on an Illumina Novaseq 6000 platform. After sequencing, we demultiplexed the reads using process_radtags from the Stacks 2.0 pipeline (47) resulting in an average of 6.31 million reads retained per sample. We assembled reads, including three outgroup samples of *L. tenuis* and *L. pedunculatus* from Durka et al. (2025) and called SNPs using dDocent 2.7.8 (48, 49) with default parameters, except of using 0.90 as ClusteringSimilarity, filtered according to O’Leary et al. 2018 introducing a mean depth threshold of 5, and a minor allele frequency threshold of 0.05 and keeping only 1 SNP per contig. We removed samples not used in this study from the resulting VCF file and re-filtered the remaining samples. The curated VCF file was converted into a fasta file (containing only the SNPs for each sample) with PGDSpider version 2.1.1.5 (50). To construct a Maximum Likelihood tree we used the program IQ-TREE release 2.0.3 (51). We used ModelFinder to select the best-fitting substitution model based on the Bayesian Information Criterion (BIC) (52) which identified the General Time Reversible (GTR) model with empirical base frequencies (F), a proportion of invariable sites (I), and gamma-distributed rate heterogeneity with four categories (G4) as the best-fit model (GTR+F+I+G4). To assess branch support, we also performed ultrafast bootstrap approximation (UFBoot) with 1,000 replicates (53). Finally, we used TreeCluster version 1.0.4 (54) to classify the SNP-based genotypes into a more restricted number of genetic groups (later used in our analyses), based on the total branch length of all leaves in the cluster (Sum Branch) with a threshold of 3.1. We visualized the phylogenetic tree using Interactive Tree of Life (iTOL) (55) and colored the samples by their genetic group assignments. From here on we will refer to these nine genetic groups as genotypes.

### Microbiome sequencing

To prepare the plant samples for microbiome sequencing, we first homogenized all frozen samples of soil, roots, shoots, flowers, and seeds with a Precellys 24 Tissue Homogenizer (Bertin Technologies, Montigny-le-Bretonneux, France), and then extracted the DNA using the FastDNA Spin Kit for Soil (MP Biomedicals, Irvine, CA, USA). We then performed a two-step PCR targeting the bacterial 16S rRNA V5-V7 region, the fungal ITS2 region, and the eukaryotic 18S rRNA V9 region, using the primers 799F/1192R, fITS7/ITS4, and F1422/R1797, respectively (Tab. S2) (20). As controls we used blank samples (UPW and blank DNA extraction). To reduce the amplification of mitochondrial and chloroplast rRNA sequences from *L. corniculatus*, we incorporated blocking oligos designed with the R package *AmpStop* (Tab. S2) (56, 57). We pooled the amplified products in equal concentrations, purified them with magnetic bead clean-up, and randomly assigned each to one of eight sequencing batches. We sequenced all pools on a Illumina MiSeq platform with PhiX control, using the MiSeq Reagent Kit v3 (600-cycle), and then processed all microbial 16S rRNA, ITS2, and 18S rRNA amplicon sequences using Mothur (56, 58, 59). To remove the primer sequences from 16S rRNA and 18S rRNA data we used Cutadapt, and to remove it from ITS2 data we used ITSx (60, 61). For the taxonomic classification of the bacterial 16S rRNA reads, we used the Greengenes database (13_8_99 release), for fungal ITS2 reads the UNITE database (02.02.2019 release), and for eukaryotic 18S rRNA the PR2 database (version 4.12.0), all of which included the PhiX genome (62–64).

### Data analysis

We carried out all data analyses in R Studio 2024.12.1 (65), and we used *dplyr* for structuring and manipulating data frames (66) *ggplot2* package (67) for creating figures. We used the *vegan* package (68) for calculating the alpha diversity (Shannon index) of bacterial, fungal or eukaryotic microbiomes of each organ in each plant, and then linear models to test for the effects of plant age or genotype. To account for variation in other factors, all models included the plant organ, and its respective interaction with plant genotype or plant age as well as sample site and year.

In addition to variation in microbial alpha diversity, we also analyzed the compositional turnover (beta diversity) between different samples, using the R packages *vegan*, *phyloseq*, and *microbiome* (68–70). We first used the OTU relative abundances for a Principal Coordinate Analysis (PCoA) of the Bray-Curtis dissimilarities between samples, and then a PERMANOVA to test how much of the variation in microbial community composition could be explained by plant age, plant genotype, or their interaction. We also visualized the relative abundance profiles of the ten most abundant classes of bacteria, fungi or eukaryotes for the different *L. corniculatus* genotypes or age groups, using the *microeco* package (69). To further understand the abundance patterns of the three different main groups of microbes, we also created graphs of the frequencies of different levels of commonness of OTUs, occurring in one to nine genotypes, or in one to five age groups.

To identify specific bacterial, fungal or eukaryotic taxa that were associated with particular plant genotypes or plant age groups, we used the R packages *microbiomeMarker* and *phyloseq* (70, 71). We used linear discriminant analysis effect size (LEfSe) (p-value < 0.05, LDA score ≥ 2) (72) to identify microbial genera that were significantly associated with particular *L. corniculatus* genotypes or age groups, either separately for each plant organ, or across organs.

## Results

The ddRAD sequencing identified a total of 5739 SNPs across all plant individuals. The Maximum Likelihood tree with best score (LL score = −733814, not shown) and consensus tree (LL score = −733810; Fig. S1) agree in all major branches. Both trees are robust, with most bootstrap values above 50%. Based on the branch lengths of the ML tree, the TreeCluster algorithm grouped the plants into nine genotype groups, each represented by 14 to 26 samples (Fig. 1b, Fig. S1). While six of these groups were significantly overrepresented in some sites (Fig. 1b, χ2 test, *P* < 0.001), all groups occurred across multiple sites (3-6 sites, average 4.4 sites), usually with multiple replicates per site, and the use of the genotype groups therefore allowed us to test for their microbiome associations at least partly independently from site variation. The estimated ages of the same plant individuals ranged from two to 13 years, with most plants being four to six years old (Fig. 1d). We found no significant relationship between plant age and sampling year (χ2 test, *P* = 0.076), but there was a statistical association between plant age and sampling site (χ2 test, *P* < 0.001), with an overrepresentation of the younger than four year old plants in one of the sites. Thus, plant age was only to a small extent confounded with site, allowing us to test for its (largely) independent association with microbiome.

### Microbiome diversity differs between plant genotypes and ages

Across all sampling sites, years, plant individuals and plant organs (672 samples), the metagenome sequencing identified a total of 4,225 bacterial (16S rRNA), 2,027 fungal (ITS2 rRNA), and 1,773 eukaryotic (18S rRNA) OTUs, clustered at 97% sequence similarity and classified into 113 phyla and 1,542 genera. The most abundant bacterial classes we detected in the *Lotus* plants were Gammaproteobacteria, Alphaproteobacteria, Betaproteobacteria, Actinobacteria, and Bacilli (Fig. 3a). The most abundant fungal classes are, Dothideomycetes, Leotiomycetes, Eurotiomycetes, Sordariomycetes, and the most abundant eukaryotic classes Insecta, Dothideomycetes, Chromadorea_X, and Agaricomycetes (Fig. 3b,c).

Part of the observed microbial diversity differed systematically between plant organs, genotypes and age groups. Specifically, the variation in Shannon diversity (= within-sample) was highly significant between the different plant organs of all genotypes and age groups. In all cases, root diversity was substantially higher than that of other organs, followed by shoot and flower diversity. Notably, seed diversity was higher than that of flowers (Fig. 2a-f, Fig. S2). Both bacterial and fungal communities differed significantly among plant genotypes, with some genotypes harboring consistently higher bacterial or fungal diversity than others, across plant organs (Fig. 2a,b). We also found a significant interaction between genotype and plant organ for bacterial diversity: genotype differences were much more pronounced in *Lotus* flowers and seeds than in shoots and roots (Fig. 2a). A similar, albeit not statistically significant, pattern was visible for fungal diversity, where also the genotype differences were greatest in flowers (Fig. 2b). In contrast to the flower and seed microbiome, the microbial diversity of roots and shoots was much less variable between plant genotypes, and there were also no genotype differences in the diversity of eukaryotes (Fig. 2c). When comparing the microbiome diversity of different plant age groups, we found a significant age effect for fungal diversity, with younger plants harboring a higher diversity of fungal microbes than older ones (Fig. 2e). However, there were no plant age effects on bacterial or eukaryote diversity (Fig. 2d,f).

**Figure 2:**
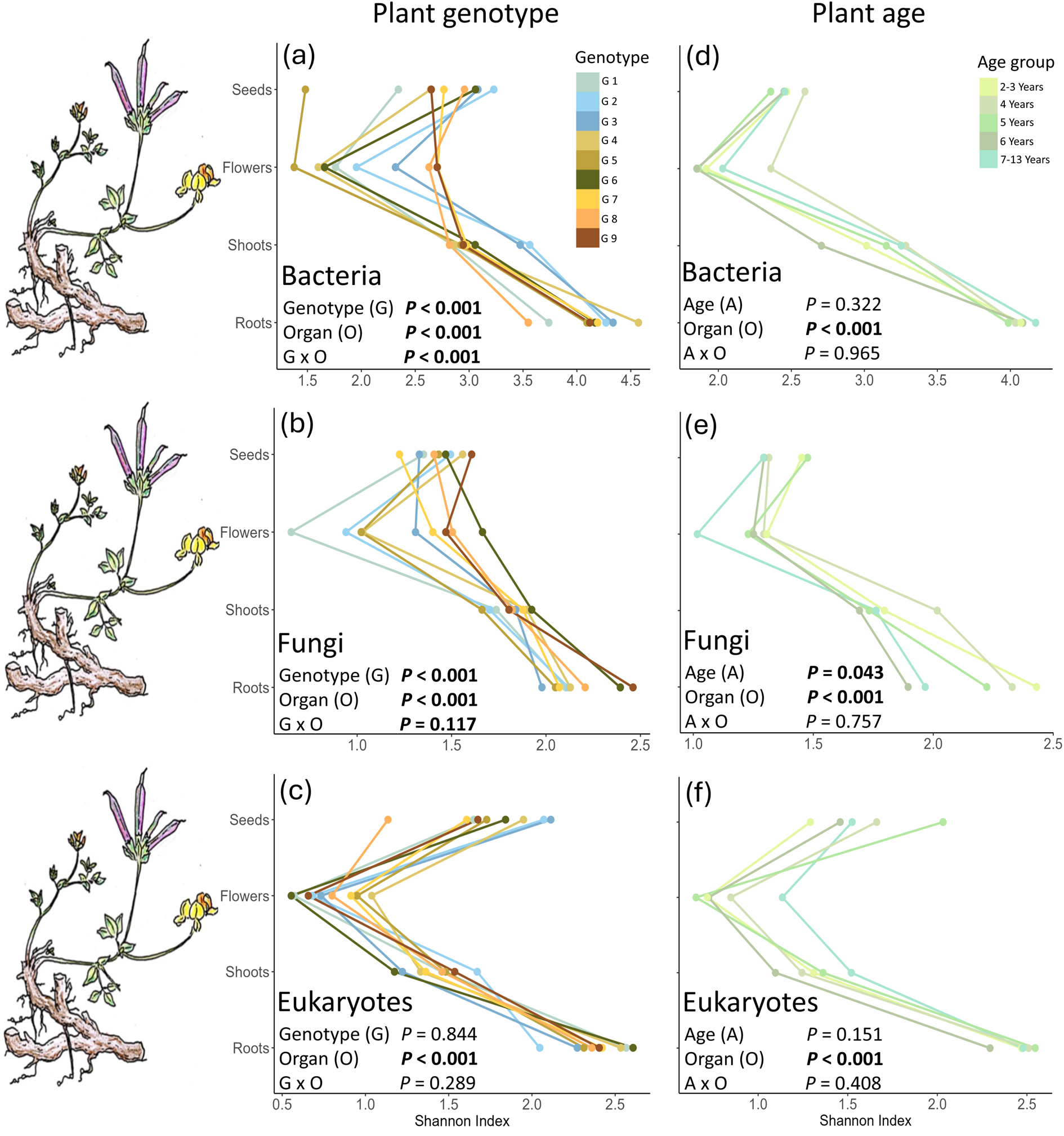
The average alpha diversity of bacterial, fungal and eukaryotic microbial communities associated with different genotypes (panels a - c) or different age groups (panels d - f) of *Lotus corniculatus* across seven semi-natural grasslands. Illustrations by S. Nicolai Rühl.

### Plant genotype and age explain variation in microbiome composition

The age and genotypes of plants were not only associated with the diversity but also the composition of their microbiome. A large number of taxa associated with *L. corniculatus* were shared among all genotypes and age groups, but we also found taxa that were less widely distributed, or even occurred in only one genotype or age group. Interestingly, the patterns of microbial commonness and rarity strongly differed between bacteria on the one side, and fungi and eukaryotes on the other side. In bacteria the majority of OTUs (2572 out of 4225) occurred in all nine genotypes, while OTUs that occurred in only one or few genotypes were rare. In fungi and eukaryotes, such less widely distributed taxa were much more frequent (Fig. 3g). The results were similar when we examined OTU distribution across age groups: almost 80% of all bacterial OTUs occurred in all age groups, and such that occurred in only one or few age groups were rare, whereas the patterns were much more even in fungi and eukaryotes (Fig. 3h).

**Figure 3:**
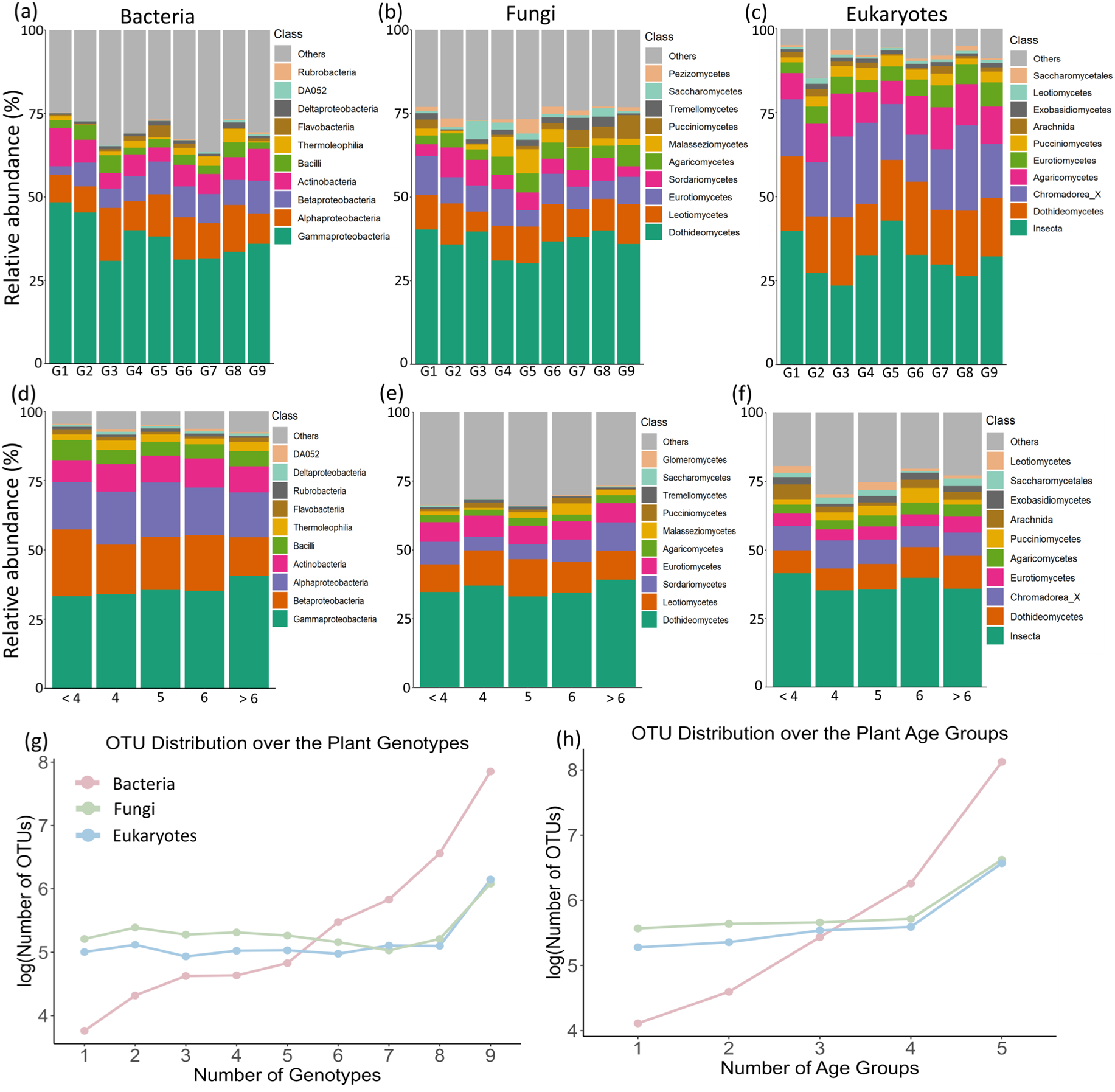
Relative abundance profiles of microbial communities associated with different genotypes (G1 - G9; panels a-c) or ages (panels d-f) of *Lotus corniculatus*, separately for bacteria, fungi and eukaryotes. We restricted the plots to the ten most abundant classes in each microbial group. Panels (g) and (h) summarize the frequencies of different levels of commonness across genotypes or ages in the three microbial groups.

The most consistent genotype effects were in roots and shoots, where the PERMANOVA identified significant plant genotype effects in the composition of all three microbial groups, with around 10% of the variation in community composition explained by plant genotype. In addition, there were significant genotype effects for bacterial communities in flowers and seeds, and for fungal communities in flowers (Tab. 1, Fig. S3). Interestingly, these genotype effects in flowers and seeds were even stronger, with over 15% variance explained.

**Table 1:** Results of PERMANOVA analyses testing the effects of plant genotype, plant age, or their interaction, on the Bray-Curtis dissimilarities of microbial communities associated with the roots, shoots, flowers, and seeds of *Lotus corniculatus* across seven semi-natural grasslands in SW Germany.

|  | Bacteria |  |  |  |  |  |  |  |  |  |  |  |
| --- | --- | --- | --- | --- | --- | --- | --- | --- | --- | --- | --- | --- |
|  | Roots |  |  | Shoots |  |  | Flowers |  |  | Seeds |  |  |
| | $R^2$ | $F$ | $Pr(>F)$ | $R^2$ | $F$ | $Pr(>F)$ | $R^2$ | $F$ | $Pr(>F)$ | $R^2$ | $F$ | $Pr(>F)$ |
| Plant genotype | 0.102 | 1.53 | **0.009** | 0.103 | 1.57 | **0.004** | 0.174 | 2.67 | **0.001** | 0.160 | 2.59 | **0.001** |
| Plant age | 0.033 | 0.99 | 0.455 | 0.043 | 1.3 | 0.107 | 0.026 | 0.81 | 0.732 | 0.049 | 1.55 | 0.064 |
| Genotype x Age | 0.234 | 1.04 | 0.309 | 0.231 | 1.04 | 0.277 | 0.180 | 0.82 | 0.956 | 0.204 | 0.98 | 0.549 |
| Residual | 0.632 |  |  | 0.623 |  |  | 0.620 |  |  | 0.588 |  |  |

|  | Fungi |  |  |  |  |  |  |  |  |  |  |  |
| --- | --- | --- | --- | --- | --- | --- | --- | --- | --- | --- | --- | --- |
|  | Roots |  |  | Shoots |  |  | Flowers |  |  | Seeds |  |  |
| | $R^2$ | $F$ | $Pr(>F)$ | $R^2$ | $F$ | $Pr(>F)$ | $R^2$ | $F$ | $Pr(>F)$ | $R^2$ | $F$ | $Pr(>F)$ |
| Plant genotype | 0.092 | 1.41 | **0.005** | 0.108 | 1.61 | **0.001** | 0.173 | 3.02 | **0.001** | 0.070 | 1.04 | 0.385 |
| Plant age | 0.029 | 0.90 | 0.71 | 0.034 | 1.06 | 0.351 | 0.042 | 1.48 | 0.103 | 0.025 | 0.74 | 0.844 |
| Genotype x Age | 0.264 | 1.21 | **0.012** | 0.248 | 1.15 | **0.036** | 0.239 | 1.23 | 0.079 | 0.258 | 1.12 | 0.385 |
| Residual | 0.615 |  |  | 0.615 |  |  | 0.545 |  |  | 0.646 |  |  |

|  | Eukaryotes |  |  |  |  |  |  |  |  |  |  |  |
| --- | --- | --- | --- | --- | --- | --- | --- | --- | --- | --- | --- | --- |
|  | Roots |  |  | Shoots |  |  | Flowers |  |  | Seeds |  |  |
| | $R^2$ | $F$ | $Pr(>F)$ | $R^2$ | $F$ | $Pr(>F)$ | $R^2$ | $F$ | $Pr(>F)$ | $R^2$ | $F$ | $Pr(>F)$ |
| Plant genotype | 0.107 | 1.66 | **0.001** | 0.090 | 1.38 | **0.008** | 0.082 | 1.21 | 0.139 | 0.073 | 1.13 | 0.235 |
| Plant age | 0.035 | 1.07 | 0.264 | 0.037 | 1.13 | 0.257 | 0.029 | 0.85 | 0.641 | 0.046 | 1.41 | 0.076 |
| Genotype x Age | 0.246 | 1.13 | **0.021** | 0.245 | 1.11 | 0.067 | 0.244 | 1.07 | 0.308 | 0.264 | 1.20 | **0.043** |
| Residual | 0.612 |  |  | 0.628 |  |  | 0.645 |  |  | 0.617 |  |  |

We found no significant main effects of plant age on the composition of bacteria, fungal and eukaryotic communities in any of the four plant organs (Tab. 1). However, there were significant plant age by genotype interactions for fungal communities in roots and shoots and eukaryotic communities in roots, indicating a complex interplay between plant genotype and age in explaining microbial community composition in these cases.

### Specific microbial taxa associated with plant genotypes and age groups

While a large number of taxa are shared among the plant genotypes and age groups, the LEfSe analyses revealed that some microbe taxa were significantly associated with particular individual plant genotypes or age groups (Fig. 4). Interestingly, these patterns of differential abundance were generally stronger in bacteria and fungi than in eukaryotic microbes, and they also tended to be more frequent with regard to plant genotypes (Fig. 4a) than for plant age groups (Fig. 4b), corroborating the PERMANOVA results with more frequent plant genotype than plant age effects (Tab. 1). Moreover, the LEfSe analyses also found that some plant genotypes and age groups showed much larger numbers of specific microbial taxa associations (e.g. genotypes 2, 3, 7 and 8, and age groups 2/3 and 5) than others. When analyzed separately for each plant organ, the LEfSE results differed significantly between organs, both for plant genotypes and plant age groups (Tab. S3), indicating that the observed microbial taxa associations are to some degree organ-specific.

**Figure 4:**
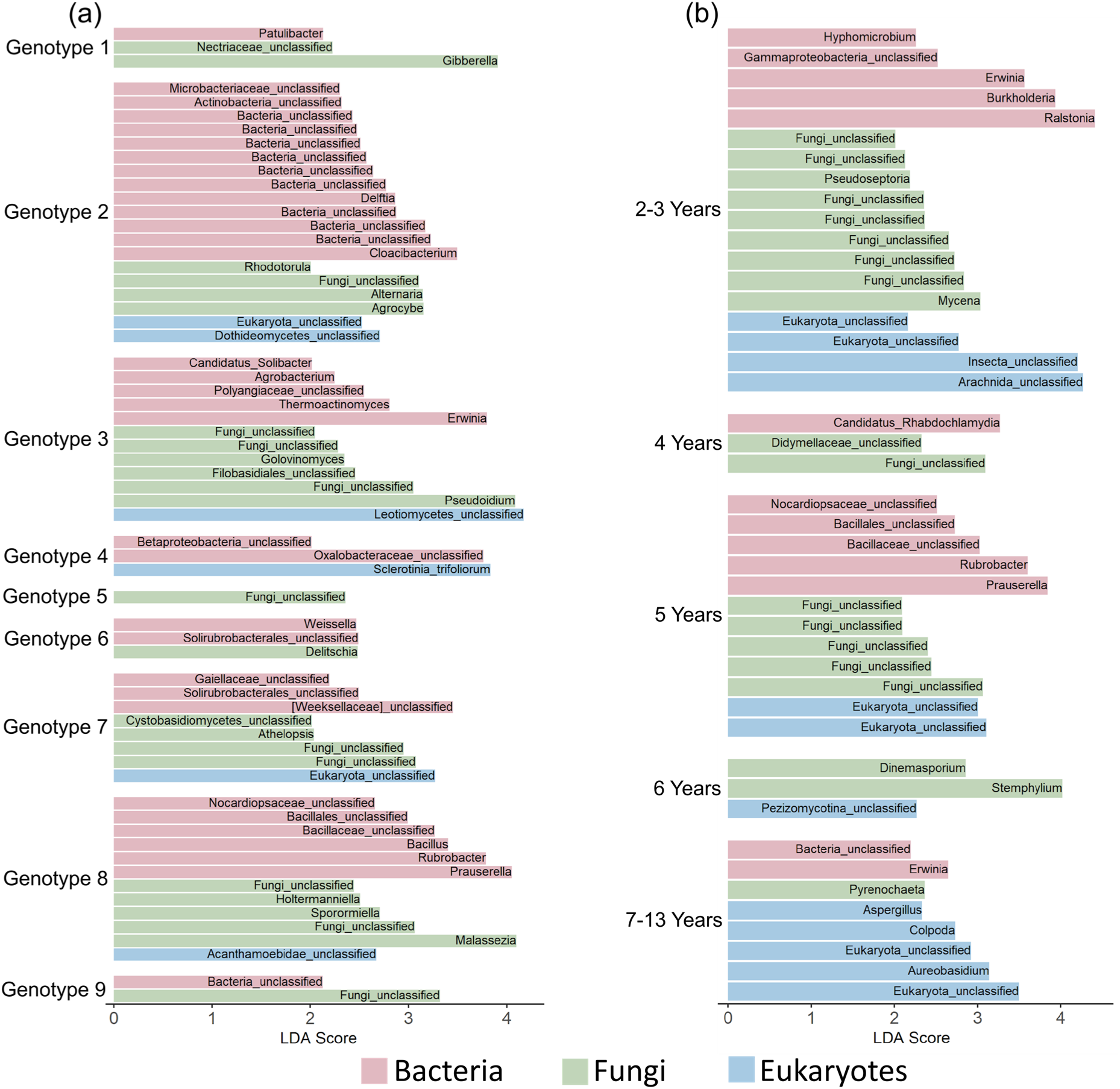
Microbial taxa significantly associated with specific genotypes (a) or age groups (b) of *Lotus corniculatus*, as identified through LefSe analyses. The three colors represent separate LefSe analyses conducted for bacteria, fungi, and eukaryotes, which were analyzed independently. These groups are presented together to provide a clearer overview of microbial variation across genotypes and age groups; however, the LDA score of taxa from different groups cannot be directly compared with each other. The raw results can be found in the supplementary material (Fig. S4).

## Discussion

Plants harbor an astonishing diversity of microbes, but the factors shaping these complex communities throughout a plant’s life, particularly in natural populations, remain poorly understood. Here, we show that in natural *Lotus corniculatus* populations both the age and genotype of plants is associated with changes in the diversity and composition of their associated microbiome. This is the case for the entire microbiome but also for individual microbial taxa that are significantly associated with specific genotypes or age groups.

### Effect of plant age and genotype on microbiome diversity

We generally found large differences in the average microbiome diversity of different plant organs, a result that has already been described in detail elsewhere (56), and that likely reflects the different properties of plant organs as habitats for microbes, and their different modes of connectedness with the environment (73). Because of this large variation, we included the plant organs as factors in our analysis, or ran analyses separately for different organs.

The alpha diversity of bacterial and fungal microorganisms differed significantly among the studied *Lotus corniculatus* genotypes. Interestingly, this effect was particularly strong in the flowers where we observed the strongest among-genotype variation in bacteria and fungi, whereas the microbiome diversity of roots and shoots was much more stable across genotypes. A possible explanation for this could be that flower traits are generally thought to be under stronger genetic control than vegetative plant traits (74), and that these traits can influence the microbiome, particularly if they affect the abundance and diversity of flower visitors (75), which are known to influence the flower microbiome (76, 77). For example, nectar secretion can vary among *L. corniculatus* genotypes (78), potentially influencing the rate of flower visitation. In the study area, the flower visitors of *L. corniculatus* include various bee species—bumblebees, honeybees, and wild bees—but also butterflies (primarily Lycaenidae) and sometimes flies and beetles (39). This broad range of pollinators may play a role in the observed flower microbiome variation between different plant genotypes. Moreover, the flower microbiome is closely linked to the seed microbiome, particularly for bacteria (79–82), which may explain why we also found considerable among-genotype variation for bacteria in seeds.

Among-genotype variation in microbiome was generally lower in plant roots and shoots than in flowers and seeds. It is well known that the root microbiome is strongly influenced, in part through affecting root traits, by different environmental factors such as soil type, structure and pH, and by the surrounding soil microbiome (1, 3, 4, 83, 84). Together, this may ‘override’ the influence of host genetics and reduce its relative impact on root microbiomes and, as a consequence, also the neighboring shoot microbiomes.

Compared to the variation among genotypes, there was less variation in microbiome diversity between plants of different ages. Only for the Shannon diversity of fungi there were significant differences between age groups. This is interesting, as many fungi play important roles as plant symbionts or pathogens (85–87), and their interactions with plants are strongly influenced by the plant immune system. The immune response of plants changes throughout their development (31, 88), and this could explain plant age-related variation in fungal microbiome diversity. Our results show that in principle plant age can also influence plant microbiome diversity, although to a lesser degree than plant genotype, and our evidence is limited to fungal microorganisms.

We should point out that comparisons of plant ages and genotypes in natural populations are not without challenges. In a field study, variation in plant age and genotype is inevitably confounded with variation in environment. We tried to account for this by aggregating both genotypes and ages into broader groups, which allowed ‘replication’ across populations. However, there was still statistical overrepresentation of some genotypes and age groups in some populations. Thus, despite our efforts both factors have not been fully independent of the sampling population, and we therefore cannot rule out that some of the observed genotype or age effects are overestimates.

### Effect of plant genotype and age on microbiome composition

We found that different *L. corniculatus* genotypes were also associated with variation in the composition of the microbiome, i.e. genetic variation in the host not only affected the alpha diversity but also the beta diversity of the microbes. Again, the extent of these effects strongly depended on the plant organ: in roots and shoots the composition of all three major microbial groups (bacteria, fungi and eukaryotes) was affected by plant genotype, whereas in flowers only the composition of bacteria and fungi, and in the seeds only that of bacteria, showed significant plant genotype associations. A plethora of genetically variable plant traits are known to affect the plant microbiome and could thus be underlying these observed composition changes. For example, root exudates are an important determinant of the rhizosphere microbiome (89, 90), and can vary across plant genotypes (91, 92). Intraspecific variation in leaf morphology and leaf chemistry has been shown to influence microbial communities in the phyllosphere (93, 94), and variation in plant defense genes is strongly affecting plant colonization by specific pathogens (95, 96).

In contrast to plant genotype, we found little evidence for effects of plant age on the composition of microbial communities. Only in fungi and eukaryotes, and mainly in roots and shoots, we found a significant plant age by genotype interaction, i.e. age effects were inconsistent and differed among genotypes, or were restricted to particular genotypes. This was counter to our expectation of age-related changes in species composition because of gene expression and functional trait changes during plant development (88). One possible reason for our findings could be that most species composition changes happen at early plant ages, and that the main factors shaping the microbiome such as root exudates (97) stabilize after this initial phase, so that the microbial communities reach an equilibrium fairly early. In our study, we deliberately collected only individuals, which had already formed inflorescences to study flowers and seeds. Thus, our youngest sampled plants were two years old, and we could not analyze the early-stage dynamics in more detail. Clearly, a better understanding of these questions, and of the observed plant age by genotype interactions, requires controlled experiments with longer-term observation of the colonization dynamics of different plant genotypes under standardized environmental conditions.

When looking at the overall frequency patterns of microbes, we found that although many taxa were shared across all plant genotypes and ages, there were still significant numbers of taxa that occurred only in a subset of plant genotypes or age groups, and there was an intriguing difference between bacteria on the one hand, and fungi and eukaryotes on the other hand. While in bacteria the level of generalism was particularly high, i.e. a large fraction of taxa occurred everywhere, frequency distributions of taxa were much more even in fungi and eukaryotes, showing a greater dispersal rates of bacteria than of non-bacteria microbes. Bacteria generally possess dispersal mechanisms unavailable to other microorganisms, allowing them to spread more efficiently. For instance, they can rapidly disperse through fungal hyphae in soil (98) and, in general, exhibit much higher dispersal rates than fungal microbes (99). In contrast, fungal dispersal is more strongly constrained by potential dispersal limitation (100), which may further contribute to the broader distribution of bacterial taxa.

### Associations of microbial taxa with specific plant genotypes and age groups

The differential abundance analysis identified a number of significant associations of microbial taxa with specific plant genotypes. Interestingly, these associations were not randomly distributed, but some plant genotypes harbored many more specific microbial taxa than others. Plant genotypes can differ in their production of secondary metabolites, which influence the recruitment or inhibition of specific microbes (101). Additionally, modifications in root morphology can limit microbial attachment and interaction, reducing the presence of specialized microbial associations in the root (102). Changes in these chemical and morphological traits can require greater levels of specialization of associated microbes.

If we look at the genotype associations in more detail, we find that several mutualistic bacteria, such as *Bacillus* (103–105), primarily soil-associated bacteria like *Cloacibacterium*, *Prauserella* and *Rubrobacter*, and some growth-promoting fungi like *Agrocybe* (106, 107) were overrepresented in some plant genotypes. Similar host genotype effects on *Bacillus* have been observed in other studies, e.g. in maize cultivars resistant to corn stalk rot versus non-resistant cultivars (108), or in sweet pepper cultivars (104). While plant interactions with the soil-associated bacteria are not yet fully understood, previous studies have shown that the plant genotype can influence the abundance of specific soil microbes in the rhizosphere (109), and that plants actively recruit genotype-specific beneficial soil microbes (110). We also found several pathogens with differential abundance among plant genotypes, e.g. *Erwinia* bacteria responsible for soft rot (111) and the fungal pathogens *Sclerotinia*, *Alternaria*, *Pseudoidium*, *Golovinomyces* and *Gibberella*. Our results corroborate previous studies with different plant species that also found genetic variation in plant resistance to these pathogens (112–115). In general, plant genetic effects on pathogen resistance are a well known phenomenon, and a key topic in agriculture and crop management (116). They also play an important role in natural populations, and are a cornerstones of the geographic mosaic of coevolution observed in many wild species (117–119).

We also found that some microbial taxa were significantly overrepresented in particular plant age groups, e.g. bacteria from the genera *Burkholderia, Erwinia and Ralstonia* and fungi from the genus *Mycena* in the youngest plant group. *Burkholderia* bacteria are known for their growth-promoting effects (110) and have been shown to benefit maize plants during germination and seedling development (120). Although the studied *Lotus* plants were already beyond these initial stages of development, the observed higher abundances of *Burkholderia* in young plants suggest that these bacteria also play a role in early *Lotus* growth and development. *Mycena* is a saprotrophic fungus opportunistically invading plant roots (121), and *Erwinia* and *Ralstonia* are bacterial pathogens causing wilt in various plant species. The increased abundances of these taxa in young *Lotus* plants suggests these may have a less well developed immune system (122, 123) or other, e.g. root physiological, differences that make them more susceptible to invasion (124). There were also microbial taxa that were particularly abundant in older plants, e.g. the pathogenic fungus *Pyrenochaeta* which is known to cause root lesions (125), and which we found overrepresented in plants older than six years. Our results confirm a previous study with oilseed rape that also observed a higher abundance of *Pyrenochaeta* in later plant developmental stages (126). In general, our data show that while plant age does not have strong effects on the overall composition of microbes, there are some significant influences at the level of individual microbial taxa, even in a long-lived plant such as *Lotus corniculatus*.

## Conclusion

Our findings show the host genotype is an important driver of natural microbiome diversity of *L. corniculatus*. This influence is evident not only in overall microbial diversity and composition but also in the abundance of specific microbial genera across different genotypes. We also found evidence of plant age influencing the fungal diversity. Additionally, the abundance of specific microbial genera varies between the different host age groups. With varied genetics or age, the physiology and requirements of the plants change, and due to the close interaction between the microbiome and the host, this can result in an altered optimal microbiome composition. Therefore, it is valuable to combine insights from controlled laboratory experiments with data from natural environments to gain a more comprehensive understanding of the impact of genotype and age on the microbiome. Overall, our results show that genotype, and to a lesser extent age, play important roles in shaping the microbiome diversity and composition of natural *L. corniculatus* populations. This knowledge can also help optimize plant health, improve crop yields, and enhance ecosystem resilience which is crucial for effective ecosystem management and plant breeding practices.

## Acknowledgements

We thank the Kemen and Bossdorf labs for help with the field sampling, in particular Elke Klenk for her invaluable assistance in collecting and processing the samples, as well Sabine Silberhorn and Julia Rafalski for their help in the lab. We are also indebted to Ralf Lauterbach and Jörg Hailer from the Schwäbische Alb local management team of the Biodiversity Exploratories for instructions and support during the field work. We have been lucky to be granted access to the Schwäbische Alb plots of the Biodiversity Exploratories (DFG Priority Program 1374) through a cooperation agreement with this project. We thank the managers of the Schwäbische Alb exploratory, Kirsten Reichel-Jung and Julia Bass and all former managers for their work in maintaining these plots and the project infrastructure, Victoria Grießmeier for her support through the central office, Andreas Ostrowski for managing the central database, and Markus Fischer, Eduard Linsenmair, Dominik Hessenmöller, Daniel Prati, Ingo Schöning, François Buscot, Ernst-Detlef Schulze, Wolfgang W. Weisser and the late Elisabeth Kalko for their role in setting up the Biodiversity Exploratories project. We thank the administration of the UNESCO Biosphere Reserve Swabian Alb as well as all landowners for the excellent collaboration. Field work permits were issued by the responsible state environmental office of Baden-Württemberg.

## Author contributions

OB, EK, FR and KL designed the study; FR and KL planned and conducted the field sampling, lab work and data analyses, JHJ helpt with the data analysis; WD helped with the library preparation and bioinformatics of the ddRAD sequencing; CR and AR helped with the age determination; FR, KL and OB wrote the manuscript with input from all authors. All authors read and approved the final manuscript.

## Competing interests

We declare that there are no conflicts of interest related to this study.

## Funding

This project has been funded by the Deutsche Forschungsgemeinschaft (DFG; projects BO 3241/11-1 to OB) as part of the priority programme 2125 Deconstruction and Reconstruction of the Plant Microbiota “DECRyPT”.

## Data availability

This work is based on data elaborated by a project associated with the Biodiversity Exploratories program (DFG Priority Program 1374). The datasets are publicly available in the Biodiversity Exploratories Information System (http://doi.org/10.17616/R32P9Q).

Sequencing data, metadata, OTU tables, and scripts are available on Biodiversity Exploratories Information System (BExIS) (https://www.bexis.uni-jena.de/) under Dataset ID 31836. Primers and blocking oligos used in this study are listed on Table S3.

## Notes

### Competing Interest Statement

The authors have declared no competing interest.

### Summary of Updates

Manuscript has been shortened, the section on the experimental test of specific lefse results has been removed (Figure 5); author list updated; citation style changed; the supplemental file updated.

