## Supplementary material for "Plant age and genotype explain variation in the microbiome of natural *Lotus corniculatus* populations"

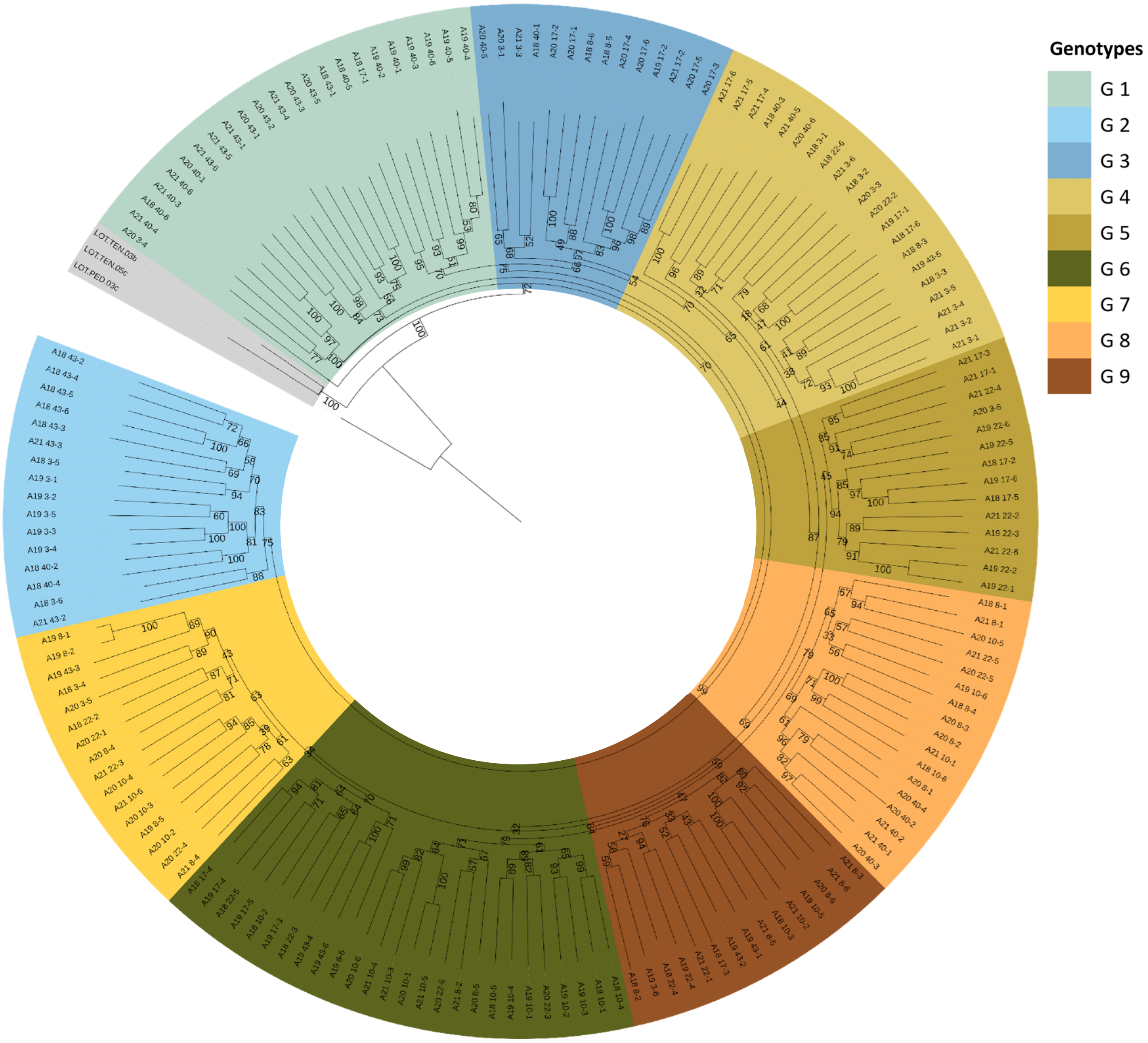


**Figure S1:** Bootstrap consensus tree derived from 1,000 replicates using IQ-TREE. Branches are supported if present in ≥50% of bootstrap replicates. Coloured by the different genotypes identified using IQ-tree.

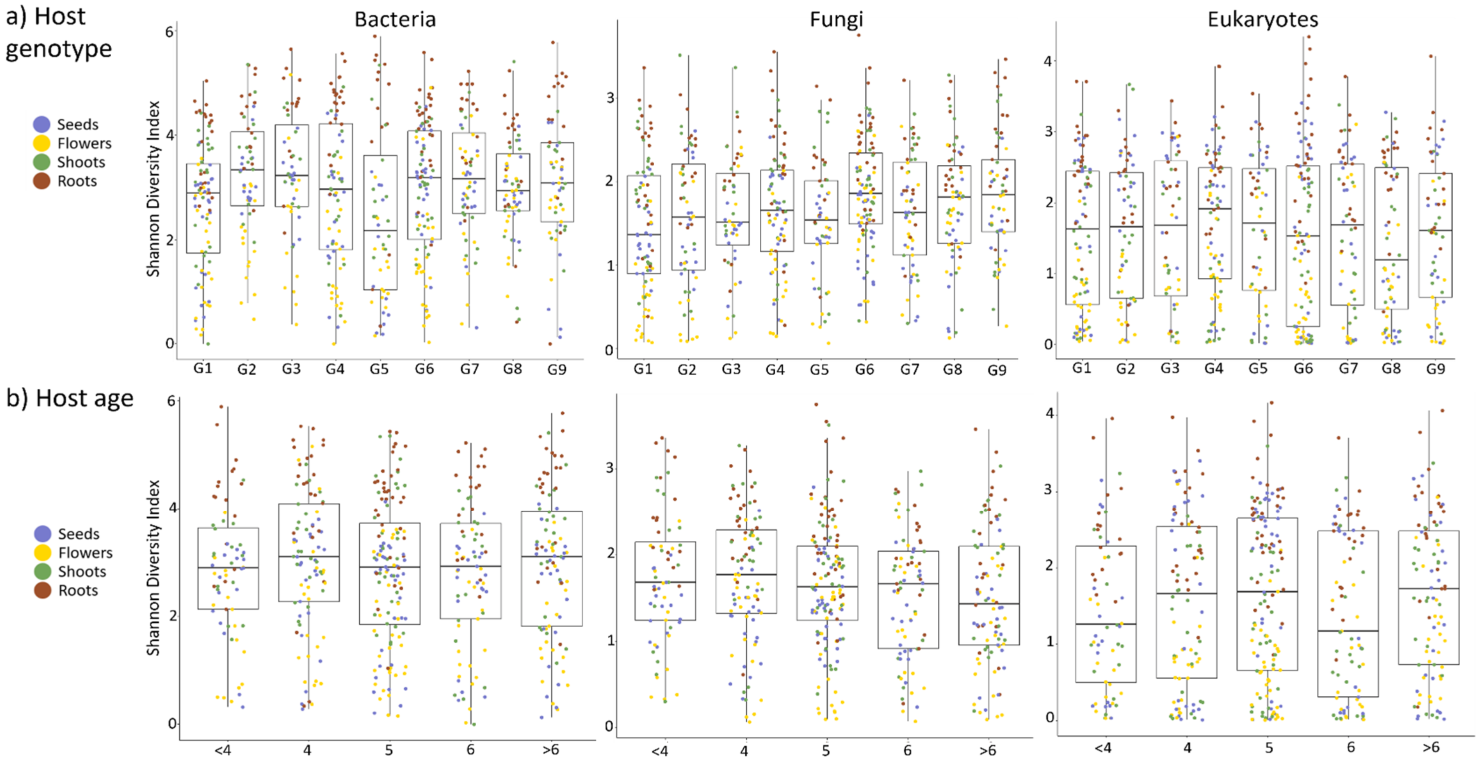


**Figure S2: a)** Box plots of the alpha diversity of bacterial, fungal and eukaryotic microbial communities associated the nine host genotypes coloured by plant organs. **b)** Box plots of the alpha diversity of bacterial, fungal and eukaryotic microbial communities associated the five host age groups coloured by plant organs.

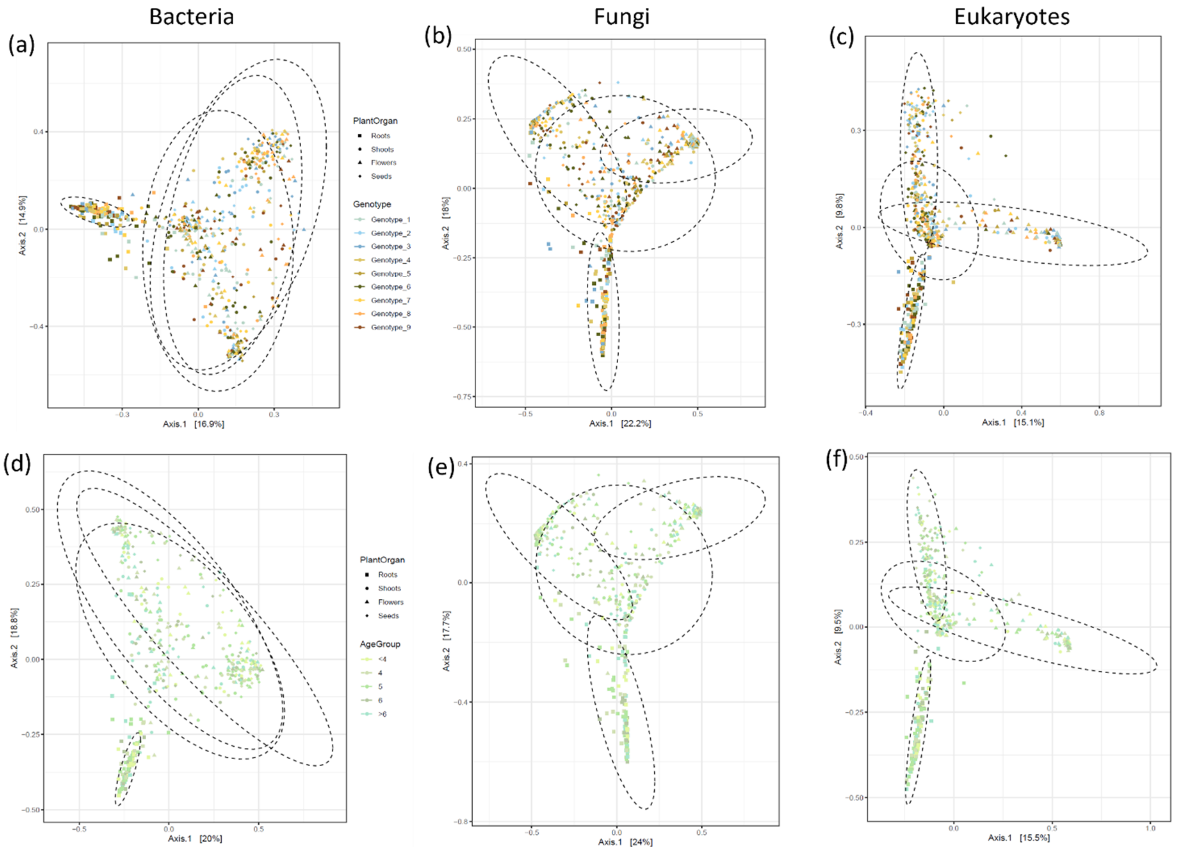


**Figure S3:** PCoA based on Bray-Curtis dissimilarities of the bacterial **(a,d),** fungal **(b,e)** and eukaryotic **(e,f)** communities associated the different plant genotypes and age groups.

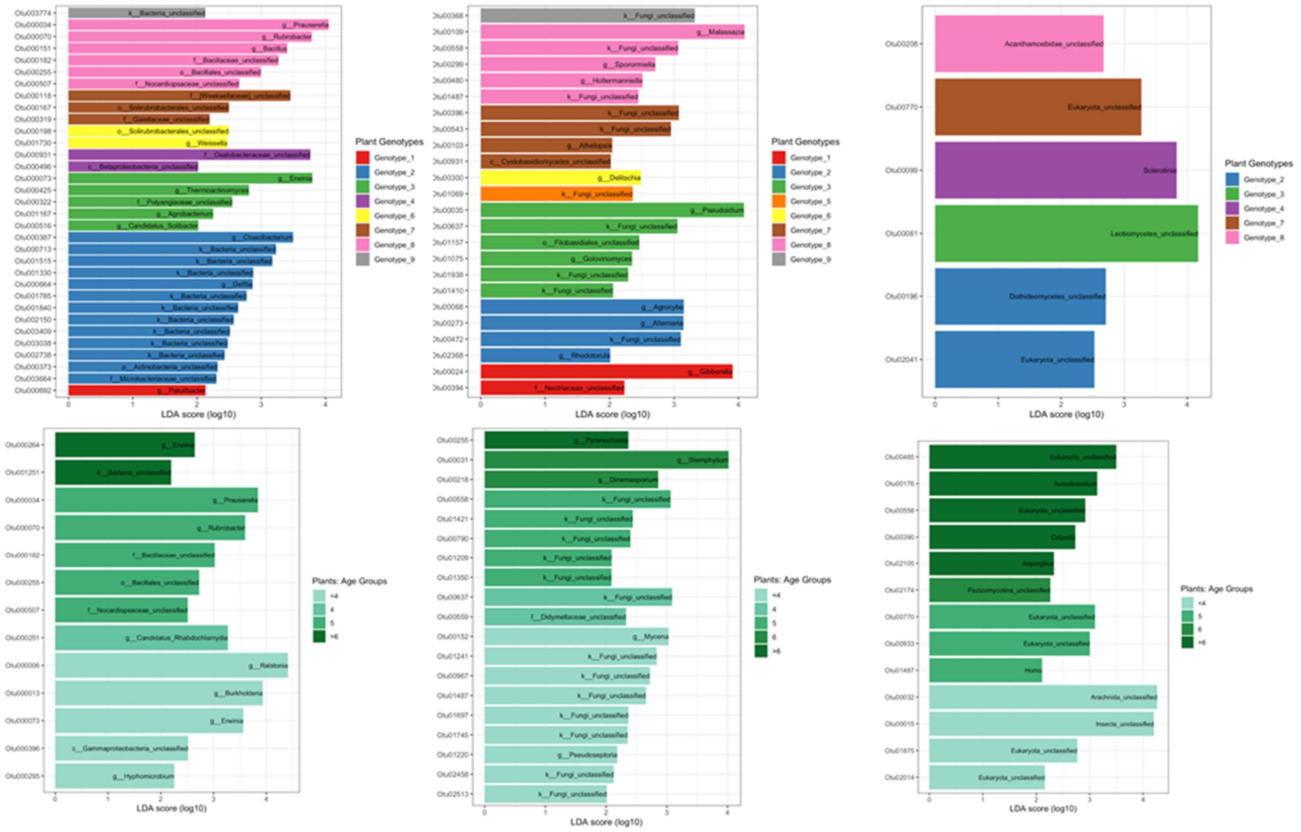


**Figure S4:** LefSe to identify differentially abundant OTUs in different genotypes of *Lotus corniculatus* plants

**Table S1:** Tab of all Age determination results including Quality score

| Sampled 2019 | | | Sampled 2020 | | | Sampled 2021 | | |
|---|---|---|---|---|---|---|---|---|
| Plant-ID | Age | Quality score | Plant-ID | Age | Quality score | Plant-ID | Age | Quality score |
| Lot19_3-1 | 4 | 2 | Lot20_3-1 | 4 | 1 | Lot21_3-1 | 5 | 2 |
| Lot19_3-2 | 5 | 3 | Lot20_3-2 | 5 | 2 | Lot21_3-2 | 3 | 2 |
| Lot19_3-3 | 5 | 2 | Lot20_3-3 | 5 | 2 | Lot21_3-3 | 4 | 1 |
| Lot19_3-4 | 7 | 3 | Lot20_3-4 | 4 | 3 | Lot21_3-4 | 3 | 1 |
| Lot19_3-5 | 11 | 2 | Lot20_3-5 | 4 | 1 | Lot21_3-5 | 4 | 1 |
| Lot19_3-6 | 4 | 2 | Lot20_3-6 | 5 | 2 | Lot21_3-6 | 8 | 2 |
| Lot19_8-1 | 4 | 2 | Lot20_8-1 | 3 | 3 | Lot21_8-1 | 4 | 3 |
| Lot19_8-2 | 8 | 1 | Lot20_8-2 | 2 | 2 | Lot21_8-2 | 3 | 3 |
| Lot19_8-3 | N/A | 4 | Lot20_8-3 | 2 | 2 | Lot21_8-3 | 4 | 3 |
| Lot19_8-4 | 4 | 1 | Lot20_8-4 | 4 | 3 | Lot21_8-4 | 3 | 3 |
| Lot19_8-5 | N/A | 4 | Lot20_8-5 | 6 | 2 | Lot21_8-5 | 3 | 2 |
| Lot19_8-6 | 6 | 2 | Lot20_8-6 | 3 | 1 | Lot21_8-6 | 4 | 3 |
| Lot19_10-1 | 5 | 1 | Lot20_10-1 | 5 | 3 | Lot21_10-1 | 8 | 2 |
| Lot19_10-2 | 4 | 1 | Lot20_10-2 | 5 | 2 | Lot21_10-2 | 3 | 2 |
| Lot19_10-3 | N/A | 4 | Lot20_10-3 | 6 | 3 | Lot21_10-3 | 5 | 1 |
| Lot19_10-4 | 6 | 3 | Lot20_10-4 | 7 | 2 | Lot21_10-4 | 5 | 3 |
| Lot19_10-5 | 8 | 2 | Lot20_10-5 | 5 | 3 | Lot21_10-5 | 5 | 1 |
| Lot19_10-6 | 6 | 2 | Lot20_10-6 | 4 | 2 | Lot21_10-6 | 7 | 2 |
| Lot19_17-1 | 3 | 1 | Lot20_17-1 | 5 | 1 | Lot21_17-1 | 3 | 3 |
| Lot19_17-2 | 4 | 3 | Lot20_17-2 | 4 | 2 | Lot21_17-2 | 5 | 2 |
| Lot19_17-3 | 3 | 1 | Lot20_17-3 | 4 | 3 | Lot21_17-3 | 4 | 1 |
| Lot19_17-4 | N/A | 4 | Lot20_17-4 | 5 | 3 | Lot21_17-4 | 6 | 1 |
| Lot19_17-5 | 8 | 3 | Lot20_17-5 | 5 | 2 | Lot21_17-5 | 6 | 1 |
| Lot19_17-6 | 6 | 2 | Lot20_17-6 | 5 | 1 | Lot21_17-6 | 5 | 2 |
| Lot19_22-1 | 5 | 1 | Lot20_22-1 | 5 | 1 | Lot21_22-1 | 5 | 2 |
| Lot19_22-2 | 5 | 2 | Lot20_22-2 | 5 | 2 | Lot21_22-2 | 5 | 2 |
| Lot19_22-3 | 5 | 3 | Lot20_22-3 | 6 | 2 | Lot21_22-3 | 6 | 3 |
| Lot19_22-4 | 8 | 3 | Lot20_22-4 | 3 | 2 | Lot21_22-4 | 4 | 2 |
| Lot19_22-5 | 5 | 2 | Lot20_22-5 | 6 | 1 | Lot21_22-5 | 4 | 1 |
| Lot19_22-6 | 6 | 2 | Lot20_22-6 | 5 | 2 | Lot21_22-6 | 4 | 3 |
| Lot19_40-1 | 6 | 3 | Lot20_40-1 | 3 | 1 | Lot21_40-1 | 6 | 1 |
| Lot19_40-2 | 10 | 3 | Lot20_40-2 | 7 | 3 | Lot21_40-2 | 8 | 2 |
| Lot19_40-3 | 12 | 3 | Lot20_40-3 | 5 | 1 | Lot21_40-3 | N/A | 4 |
| Lot19_40-4 | 10 | 3 | Lot20_40-4 | 6 | 3 | Lot21_40-4 | 7 | 2 |
| Lot19_40-5 | 8 | 2 | Lot20_40-5 | 7 | 2 | Lot21_40-5 | N/A | 4 |
| Lot19_40-6 | 6 | 2 | Lot20_40-6 | 6 | 2 | Lot21_40-6 | 3 | 2 |
| Lot19_43-1 | 8 | 3 | Lot20_43-1 | 6 | 3 | Lot21_43-1 | 8 | 1 |
| Lot19_43-2 | 8 | 2 | Lot20_43-2 | 6 | 1 | Lot21_43-2 | 3 | 1 |
| Lot19_43-3 | 6 | 2 | Lot20_43-3 | 5 | 3 | Lot21_43-3 | 5 | 1 |
| Lot19_43-4 | 4 | 2 | Lot20_43-4 | 5 | 3 | Lot21_43-4 | 10 | 2 |
| Lot19_43-5 | 13 | 2 | Lot20_43-5 | 7 | 2 | Lot21_43-5 | 5 | 2 |
| Lot19_43-6 | 5 | 2 | Lot20_43-6 | 4 | 2 | Lot21_43-6 | 5 | 2 |

**Table S2:** Primers and blocking oligos used for the microbiome sequencing.

| Primer name | Primer sequence (5'-to-3' orientation) |
|---|---|
| 799F | AACMGGATTAGATACCCKG |
| 1192R | ACGTCATCCCCACCTTCC |
| fITS7 | GTGARTCATCGAATCTTTG |
| ITS4 | TCCTCCGCTTATTGATATGC |
| F1422 | ATAACAGGTCTGTGATGCCC |
| R1797 | TGATCCTTCTGCAGGTTCACCTAC |
| clamp1_BV5_mitoF | GATGAGTGTTCGCCCTTGGTCTACGTGGAT |
| clamp1_BV5_mitoR | CTGCTCAGGGTTCCAAACTCAACGTTGGCA |
| clamp1_ITS2_F | AACCATTAGGTCGAGGGCACGTCTGCCTGG |
| clamp1_ITS2_R | TGAGMGYGGTTACACCACGCATGCGGGTCT |
| clamp9_PV9_F | GATGTATTCAACGAGTCTATAGCCTTGGCC |
| clamp15_PV9_R | TCTCACAACGTCGCAGGCAGCGAACCGCCC |

**Table S3:** LefSe to identify differentially abundant OTUs in different genotypes of *Lotus corniculatus* plants by plant organs.

| **Genotype** | | | | | | | |
|---|---|---|---|---|---|---|---|
| **OTU** | **enrich_genotype** | **organ** | **ef_lda** | **pvalue** | **padj** | **Locus** | **Genus** |
| Otu000030 | Genotype_4 | roots | 3.589 | <0.001 | <0.001 | 16S | o__0319-7L14_unclassified |
| Otu000074 | Genotype_8 | roots | 3.739 | 0.003 | 0.003 | 16S | f__Sinobacteraceae_unclassified |
| Otu000075 | Genotype_3 | roots | 3.346 | <0.001 | <0.001 | 16S | f__Solirubrobacteraceae_unclassified |
| Otu000098 | Genotype_3 | roots | 3.184 | <0.001 | <0.001 | 16S | g__Mycobacterium |
| Otu000103 | Genotype_4 | roots | 3.052 | 0.004 | 0.004 | 16S | g__Kribbella |
| Otu000107 | Genotype_1 | roots | 3.638 | 0.037 | 0.037 | 16S | f__Sphingomonadaceae_unclassified |
| Otu000110 | Genotype_7 | roots | 3.423 | 0.000 | 0.000 | 16S | o__Solirubrobacterales_unclassified |
| Otu000129 | Genotype_4 | roots | 2.890 | <0.001 | <0.001 | 16S | f__Gaiellaceae_unclassified |
| Otu000133 | Genotype_1 | roots | 3.399 | 0.001 | 0.001 | 16S | g__Agrobacterium |
| Otu000162 | Genotype_5 | roots | 2.747 | 0.001 | 0.001 | 16S | f__[Entotheonellaceae]_unclassified |
| Otu000167 | Genotype_6 | roots | 2.895 | 0.016 | 0.016 | 16S | o__Solirubrobacterales_unclassified |
| Otu000193 | Genotype_5 | roots | 2.737 | 0.000 | 0.000 | 16S | f__EB1017_unclassified |
| Otu000215 | Genotype_5 | roots | 2.596 | 0.002 | 0.002 | 16S | f__Gaiellaceae_unclassified |
| Otu000244 | Genotype_5 | roots | 2.659 | <0.001 | <0.001 | 16S | g__Amaricoccus |
| Otu000254 | Genotype_2 | roots | 2.981 | <0.001 | <0.001 | 16S | c_Gammaproteobacteria_unclassified |
| Otu000260 | Genotype_4 | roots | 2.683 | <0.001 | <0.001 | 16S | o__Solirubrobacterales_unclassified |
| Otu000263 | Genotype_4 | roots | 2.507 | <0.001 | <0.001 | 16S | f__C111_unclassified |
| Otu000273 | Genotype_5 | roots | 2.203 | 0.016 | 0.016 | 16S | c__PAUC37f_unclassified |
| Otu000282 | Genotype_6 | roots | 2.485 | 0.001 | 0.001 | 16S | f__Micromonosporaceae_unclassified |
| Otu000283 | Genotype_2 | roots | 2.648 | 0.008 | 0.008 | 16S | f__Gaiellaceae_unclassified |
| Otu000289 | Genotype_1 | roots | 2.539 | 0.001 | 0.001 | 16S | g__Marmoricola |
| Otu000338 | Genotype_9 | roots | 2.966 | 0.006 | 0.006 | 16S | f__Haliangiaceae_unclassified |
| Otu000351 | Genotype_4 | roots | 2.310 | <0.001 | <0.001 | 16S | o__Acidimicrobiales_unclassified |
| Otu000364 | Genotype_3 | roots | 2.408 | <0.001 | <0.001 | 16S | f__AKIW874_unclassified |
| Otu000368 | Genotype_4 | roots | 2.385 | <0.001 | <0.001 | 16S | o__Acidimicrobiales_unclassified |
| Otu000370 | Genotype_3 | roots | 2.479 | 0.001 | 0.001 | 16S | c__S085_unclassified |
| Otu000373 | Genotype_2 | roots | 2.852 | <0.001 | <0.001 | 16S | p__Actinobacteria_unclassified |

| | | | | | | | |
|---|---|---|---|---|---|---|---|
| Otu000386 | Genotype_5 | roots | 2.358 | <0.001 | <0.001 | 16S | f__EB1017_unclassified |
| Otu000423 | Genotype_5 | roots | 2.425 | <0.001 | <0.001 | 16S | g__Balneimonas |
| Otu000425 | Genotype_3 | roots | 2.514 | 0.000 | 0.000 | 16S | g__Thermoactinomyces |
| Otu000436 | Genotype_3 | roots | 2.303 | 0.002 | 0.002 | 16S | g__Coprococcus |
| Otu000467 | Genotype_3 | roots | 2.656 | 0.002 | 0.002 | 16S | f__Comamonadaceae_unclassified |
| Otu000474 | Genotype_9 | roots | 3.100 | <0.001 | <0.001 | 16S | g__Sphingomonas |
| Otu000516 | Genotype_4 | roots | 2.096 | <0.001 | <0.001 | 16S | g__Candidatus_Solibacter |
| Otu000536 | Genotype_8 | roots | 2.365 | 0.035 | 0.035 | 16S | f__Dolo_23_unclassified |
| Otu000545 | Genotype_8 | roots | 2.650 | <0.001 | <0.001 | 16S | g__Actinoallomurus |
| Otu000567 | Genotype_2 | roots | 2.038 | <0.001 | <0.001 | 16S | f__EB1017_unclassified |
| Otu000573 | Genotype_5 | roots | 2.085 | <0.001 | <0.001 | 16S | o__Acidimicrobiales_unclassified |
| Otu000592 | Genotype_9 | roots | 2.088 | 0.016 | 0.016 | 16S | f__Conexibacteraceae_unclassified |
| Otu000644 | Genotype_5 | roots | 2.362 | <0.001 | <0.001 | 16S | f__Cystobacterineae_unclassified |
| Otu000656 | Genotype_5 | roots | 2.133 | <0.001 | <0.001 | 16S | o__Micrococcales_unclassified |
| Otu000669 | Genotype_8 | roots | 2.470 | 0.006 | 0.006 | 16S | f__Caulobacteraceae_unclassified |
| Otu000690 | Genotype_7 | roots | 2.575 | 0.002 | 0.002 | 16S | c_Gammaproteobacteria_unclassified |
| Otu000734 | Genotype_5 | roots | 2.029 | 0.014 | 0.014 | 16S | o__NB1-j_unclassified |
| Otu000764 | Genotype_2 | roots | 2.011 | 0.004 | 0.004 | 16S | f__Haliangiaceae_unclassified |
| Otu000812 | Genotype_2 | roots | 2.423 | <0.001 | <0.001 | 16S | f__Polyangiaceae_unclassified |
| Otu000974 | Genotype_2 | roots | 2.077 | 0.026 | 0.026 | 16S | f__Beijerinckiaceae_unclassified |
| Otu001053 | Genotype_8 | roots | 2.559 | 0.002 | 0.002 | 16S | c__SJA-4_unclassified |
| Otu001075 | Genotype_2 | roots | 2.479 | <0.001 | <0.001 | 16S | g__Rahnella |
| Otu001147 | Genotype_3 | roots | 2.025 | <0.001 | <0.001 | 16S | o__Bacillales_unclassified |
| Otu001210 | Genotype_5 | roots | 2.091 | 0.024 | 0.024 | 16S | g__Candidatus_Protochlamydia |
| Otu00004 | Genotype_5 | roots | 5.014 | 0.027 | 0.027 | ITS2 | g__Exophiala |
| Otu00035 | Genotype_7 | roots | 2.227 | 0.007 | 0.007 | ITS2 | g__Pseudoidium |
| Otu00038 | Genotype_1 | roots | 4.110 | 0.017 | 0.017 | ITS2 | f__Bionectriaceae_unclassified |
| Otu00068 | Genotype_2 | roots | 3.920 | <0.001 | <0.001 | ITS2 | g__Agrocybe |
| Otu00139 | Genotype_3 | roots | 3.419 | 0.007 | 0.007 | ITS2 | g__Leohumicola |
| Otu00300 | Genotype_1 | roots | 3.000 | 0.013 | 0.013 | ITS2 | g__Delitschia |
| Otu00774 | Genotype_1 | roots | 2.588 | <0.001 | <0.001 | ITS2 | g__Dendryphion |
| Otu01037 | Genotype_5 | roots | 2.037 | 0.002 | 0.002 | ITS2 | g__Gremmenia |
| Otu01062 | Genotype_9 | roots | 2.006 | 0.002 | 0.002 | ITS2 | g__Microdochium |
| Otu00023 | Genotype_6 | roots | 2.829 | 0.006 | 0.006 | 18S | Penicillium |
| Otu00024 | Genotype_1 | roots | 4.438 | 0.004 | 0.004 | 18S | Chromadorea_X_unclassified |
| Otu00129 | Genotype_4 | roots | 2.948 | 0.004 | 0.004 | 18S | Gregarinidae_unclassified |
| Otu00154 | Genotype_3 | roots | 2.636 | 0.012 | 0.012 | 18S | Chytridiomycotina_unclassified |
| Otu00196 | Genotype_2 | roots | 3.218 | <0.001 | <0.001 | 18S | Dothideomycetes_unclassified |
| Otu00208 | Genotype_8 | roots | 3.401 | 0.022 | 0.022 | 18S | Acanthamoebidae_unclassified |
| Otu00306 | Genotype_6 | roots | 2.922 | 0.002 | 0.002 | 18S | Assulina |
| Otu00314 | Genotype_2 | roots | 2.355 | 0.015 | 0.015 | 18S | Trichosporon |
| Otu00505 | Genotype_6 | roots | 2.313 | 0.037 | 0.037 | 18S | Ramicandelaber |
| Otu00510 | Genotype_9 | roots | 2.552 | 0.005 | 0.005 | 18S | Eukaryota_unclassified |
| Otu00770 | Genotype_3 | roots | 3.275 | 0.020 | 0.020 | 18S | Eukaryota_unclassified |
| Otu000009 | Genotype_3 | shoots | 2.987 | 0.010 | 0.010 | 16S | g__Phyllobacterium |
| Otu000017 | Genotype_2 | shoots | 2.617 | <0.001 | <0.001 | 16S | g__Mesorhizobium |
| Otu000034 | Genotype_3 | shoots | 4.025 | 0.001 | 0.001 | 16S | g__Prauserella |
| Otu000062 | Genotype_2 | shoots | 3.000 | 0.030 | 0.030 | 16S | g__Flavobacterium |
| Otu000070 | Genotype_3 | shoots | 3.739 | 0.002 | 0.002 | 16S | g__Rubrobacter |
| Otu000073 | Genotype_3 | shoots | 3.857 | <0.001 | <0.001 | 16S | g__Erwinia |

| | | | | | | | |
|---|---|---|---|---|---|---|---|
| Otu000118 | Genotype_8 | shoots | 3.148 | 0.011 | 0.011 | 16S | f__[Weeksellaceae]_unclassified |
| Otu000151 | Genotype_3 | shoots | 3.351 | 0.001 | 0.001 | 16S | g__Bacillus |
| Otu000466 | Genotype_2 | shoots | 2.792 | 0.002 | 0.002 | 16S | g__Massilia |
| Otu000938 | Genotype_5 | shoots | 2.025 | 0.001 | 0.001 | 16S | f__Sporichthyaceae_unclassified |
| Otu001023 | Genotype_2 | shoots | 2.012 | 0.013 | 0.013 | 16S | f__Xanthomonadaceae_unclassified |
| Otu001029 | Genotype_1 | shoots | 2.250 | 0.000 | 0.000 | 16S | g__Microbacterium |
| Otu001174 | Genotype_9 | shoots | 2.080 | 0.012 | 0.012 | 16S | f__Planococcaceae_unclassified |
| Otu001270 | Genotype_1 | shoots | 2.387 | <0.001 | <0.001 | 16S | f__Nocardiaceae_unclassified |
| Otu001782 | Genotype_2 | shoots | 2.658 | 0.000 | 0.000 | 16S | g__Buchnera |
| Otu001914 | Genotype_2 | shoots | 2.352 | <0.001 | <0.001 | 16S | f__Oxalobacteraceae_unclassified |
| Otu002076 | Genotype_2 | shoots | 2.022 | <0.001 | <0.001 | 16S | f__Actinosynnemataceae_unclassified |
| Otu003094 | Genotype_6 | shoots | 2.137 | <0.001 | <0.001 | 16S | g__Pseudomonas |
| Otu00004 | Genotype_2 | shoots | 3.176 | 0.003 | 0.003 | ITS2 | g__Exophiala |
| Otu00026 | Genotype_3 | shoots | 2.411 | <0.001 | <0.001 | ITS2 | g__Mycena |
| Otu00103 | Genotype_2 | shoots | 2.376 | 0.000 | 0.000 | ITS2 | g__Athelopsis |
| Otu00122 | Genotype_2 | shoots | 3.164 | <0.001 | <0.001 | ITS2 | g__Plectosphaerella |
| Otu00312 | Genotype_7 | shoots | 3.324 | 0.006 | 0.006 | ITS2 | g__Dissoconium |
| Otu00381 | Genotype_9 | shoots | 2.886 | <0.001 | <0.001 | ITS2 | g__Cystofilobasidium |
| Otu00386 | Genotype_2 | shoots | 2.021 | 0.005 | 0.005 | ITS2 | f__Lentitheciaceae_unclassified |
| Otu00562 | Genotype_1 | shoots | 2.729 | <0.001 | <0.001 | ITS2 | o__Xylariales_unclassified |
| Otu01017 | Genotype_2 | shoots | 2.427 | 0.001 | 0.001 | ITS2 | g__Desmococcus |
| Otu01075 | Genotype_7 | shoots | 2.665 | 0.003 | 0.003 | ITS2 | g__Golovinomyces |
| Otu01359 | Genotype_5 | shoots | 2.390 | 0.000 | 0.000 | ITS2 | o__Pleosporales_unclassified |
| Otu01433 | Genotype_2 | shoots | 2.005 | 0.002 | 0.002 | ITS2 | f__Stachybotryaceae_unclassified |
| Otu01689 | Genotype_2 | shoots | 2.096 | 0.000 | 0.000 | ITS2 | g__unclassified_Verrucariaceae |
| Otu01708 | Genotype_5 | shoots | 2.227 | 0.000 | 0.000 | ITS2 | g__Puccinia |
| Otu01710 | Genotype_2 | shoots | 2.235 | 0.000 | 0.000 | ITS2 | o__Hypocreales_unclassified |
| Otu01762 | Genotype_3 | shoots | 2.072 | <0.001 | <0.001 | ITS2 | g__Thanatephorus |
| Otu02224 | Genotype_2 | shoots | 2.039 | <0.001 | <0.001 | ITS2 | g__Hypoxylon |
| Otu00043 | Genotype_2 | shoots | 3.314 | 0.032 | 0.032 | 18S | Plectus |
| Otu00308 | Genotype_9 | shoots | 3.054 | <0.001 | <0.001 | 18S | Microdochium |
| Otu02170 | Genotype_2 | shoots | 2.556 | 0.000 | 0.000 | 18S | Klebsormidium |
| Otu000016 | Genotype_8 | flowers | 4.265 | <0.001 | <0.001 | 16S | g__Burkholderia |
| Otu000024 | Genotype_3 | flowers | 2.793 | 0.043 | 0.043 | 16S | f__Xanthomonadaceae_unclassified |
| Otu000025 | Genotype_1 | flowers | 3.196 | 0.035 | 0.035 | 16S | g__Rhodoplanes |
| Otu000034 | Genotype_8 | flowers | 4.140 | <0.001 | <0.001 | 16S | g__Prauserella |
| Otu000070 | Genotype_8 | flowers | 3.843 | <0.001 | <0.001 | 16S | g__Rubrobacter |
| Otu000073 | Genotype_8 | flowers | 3.893 | <0.001 | <0.001 | 16S | g__Erwinia |
| Otu000082 | Genotype_8 | flowers | 3.730 | <0.001 | <0.001 | 16S | p__Proteobacteria_unclassified |
| Otu000086 | Genotype_8 | flowers | 3.599 | <0.001 | <0.001 | 16S | f__Burkholderiaceae_unclassified |
| Otu000096 | Genotype_7 | flowers | 2.767 | 0.009 | 0.009 | 16S | g__Bradyrhizobium |
| Otu000115 | Genotype_2 | flowers | 3.033 | 0.029 | 0.029 | 16S | g__Massilia |
| Otu000151 | Genotype_8 | flowers | 3.536 | <0.001 | <0.001 | 16S | g__Bacillus |
| Otu000291 | Genotype_8 | flowers | 2.941 | 0.004 | 0.004 | 16S | g__Methylotenera |
| Otu000404 | Genotype_6 | flowers | 2.339 | 0.003 | 0.003 | 16S | f__Sinobacteraceae_unclassified |
| Otu001113 | Genotype_2 | flowers | 2.511 | 0.001 | 0.001 | 16S | c_Gammaproteobacteria_unclassified |
| Otu001383 | Genotype_7 | flowers | 2.113 | <0.001 | <0.001 | 16S | o__MIZ46_unclassified |
| Otu004213 | Genotype_2 | flowers | 2.629 | <0.001 | <0.001 | 16S | g__Deinococcus |
| Otu00273 | Genotype_2 | flowers | 3.742 | <0.001 | <0.001 | ITS2 | g__Alternaria |
| Otu00543 | Genotype_8 | flowers | 3.115 | <0.001 | <0.001 | ITS2 | k__Fungi_unclassified |

| | | | | | | | |
|---|---|---|---|---|---|---|---|
| Otu01410 | Genotype_7 | flowers | 2.273 | <0.001 | <0.001 | ITS2 | k__Fungi_unclassified |
| Otu02368 | Genotype_2 | flowers | 2.398 | <0.001 | <0.001 | ITS2 | g__Rhodotorula |
| Otu00099 | Genotype_8 | flowers | 4.061 | 0.011 | 0.011 | 18S | Sclerotinia |
| Otu000126 | Genotype_4 | seeds | 2.395 | 0.024 | 0.024 | 16S | g__Rhizobium |
| Otu000141 | Genotype_2 | seeds | 4.070 | <0.001 | <0.001 | 16S | o__Ellin6513_unclassified |
| Otu000332 | Genotype_2 | seeds | 4.604 | 0.011 | 0.011 | 16S | k__Bacteria_unclassified |
| Otu000493 | Genotype_5 | seeds | 2.306 | <0.001 | <0.001 | 16S | g__Pantoea |
| Otu000630 | Genotype_5 | seeds | 2.115 | 0.003 | 0.003 | 16S | g__Pantoea |
| Otu000713 | Genotype_6 | seeds | 3.654 | <0.001 | <0.001 | 16S | k__Bacteria_unclassified |
| Otu001330 | Genotype_2 | seeds | 3.344 | <0.001 | <0.001 | 16S | k__Bacteria_unclassified |
| Otu001840 | Genotype_2 | seeds | 3.150 | 0.001 | 0.001 | 16S | k__Bacteria_unclassified |
| Otu002150 | Genotype_2 | seeds | 3.025 | 0.001 | 0.001 | 16S | k__Bacteria_unclassified |
| Otu00321 | Genotype_8 | seeds | 4.259 | 0.024 | 0.024 | ITS2 | g__Malassezia |
| Otu00410 | Genotype_2 | seeds | 3.392 | 0.005 | 0.005 | ITS2 | k__Fungi_unclassified |
| Otu00472 | Genotype_2 | seeds | 3.583 | <0.001 | <0.001 | ITS2 | k__Fungi_unclassified |
| Otu00603 | Genotype_2 | seeds | 3.529 | 0.009 | 0.009 | ITS2 | k__Fungi_unclassified |
| Otu00990 | Genotype_3 | seeds | 3.127 | 0.005 | 0.005 | ITS2 | k__Fungi_unclassified |

### Age

| OTU | enrich_age | organ | ef_lda | pvalue | padj | Locus | Genus |
|---|---|---|---|---|---|---|---|
| Otu000093 | >6 | roots | 3.680 | 0.002 | 0.002 | 16S | f__Xanthobacteraceae_unclassified |
| Otu000295 | <4 | roots | 2.716 | 0.015 | 0.015 | 16S | g__Hyphomicrobium |
| Otu000396 | <4 | roots | 3.184 | 0.020 | 0.020 | 16S | c__Gammaproteobacteria_unclassified |
| Otu000421 | >6 | roots | 2.634 | 0.003 | 0.003 | 16S | g__Steroidobacter |
| Otu000575 | >6 | roots | 2.282 | 0.029 | 0.029 | 16S | f__C111_unclassified |
| Otu000889 | <4 | roots | 2.184 | 0.013 | 0.013 | 16S | g__Bacillus |
| Otu001115 | <4 | roots | 2.086 | 0.046 | 0.046 | 16S | o__B07_WMSP1_unclassified |
| Otu001130 | >6 | roots | 2.007 | 0.036 | 0.036 | 16S | o__Phycisphaerales_unclassified |
| Otu001174 | <4 | roots | 2.045 | 0.048 | 0.048 | 16S | f__Planococcaceae_unclassified |
| Otu001384 | 6 | roots | 2.061 | 0.047 | 0.047 | 16S | f__Flavobacteriaceae_unclassified |
| Otu001569 | <4 | roots | 2.164 | 0.043 | 0.043 | 16S | g__Bdellovibrio |
| Otu001627 | >6 | roots | 2.164 | <0.001 | <0.001 | 16S | g__Criblamydia |
| Otu00152 | <4 | roots | 3.717 | 0.008 | 0.008 | ITS2 | g__Mycena |
| Otu00246 | 4 | roots | 2.817 | 0.032 | 0.032 | ITS2 | p__Ascomycota_unclassified |
| Otu00255 | >6 | roots | 2.964 | 0.006 | 0.006 | ITS2 | g__Pyrenochaeta |
| Otu00816 | >6 | roots | 2.285 | 0.002 | 0.002 | ITS3 | g__Rhizophagus |
| Otu00072 | 6 | roots | 4.353 | 0.018 | 0.018 | 18S | Helicotylenchus |
| Otu00223 | 4 | roots | 3.070 | 0.044 | 0.044 | 18S | Pezizomycotina_unclassified |
| Otu00413 | >6 | roots | 2.628 | 0.004 | 0.004 | 18S | Pezizomycotina_unclassified |
| Otu00469 | 5 | roots | 2.097 | 0.030 | 0.030 | 18S | Syncephalis |
| Otu00577 | <4 | roots | 2.292 | 0.037 | 0.037 | 18S | Chromadorea_X_unclassified |
| Otu01077 | <4 | roots | 2.005 | <0.001 | <0.001 | 18S | Paulinella |
| Otu000033 | >6 | shoots | 3.370 | 0.045 | 0.045 | 16S | g__Chryseobacterium |
| Otu000043 | 4 | shoots | 2.542 | 0.001 | 0.001 | 16S | f__Bradyrhizobiaceae_unclassified |
| Otu000070 | 4 | shoots | 3.672 | 0.007 | 0.007 | 16S | g__Rubrobacter |
| Otu000186 | >6 | shoots | 2.484 | 0.034 | 0.034 | 16S | c__S085_unclassified |
| Otu000191 | >6 | shoots | 2.835 | 0.046 | 0.046 | 16S | f__Gaiellaceae_unclassified |
| Otu000370 | >6 | shoots | 2.106 | 0.020 | 0.020 | 16S | c__S085_unclassified |
| Otu000511 | >6 | shoots | 2.798 | 0.013 | 0.013 | 16S | g__Sphingomonas |
| Otu000560 | >6 | shoots | 2.203 | 0.030 | 0.030 | 16S | g__Nocardioides |
| Otu000568 | 4 | shoots | 2.360 | 0.028 | 0.028 | 16S | g__Streptomyces |

| | | | | | | | |
|---|---|---|---|---|---|---|---|
| Otu000977 | >6 | shoots | 2.130 | 0.013 | 0.013 | 16S | f__Gaiellaceae_unclassified |
| Otu001155 | <4 | shoots | 2.117 | 0.040 | 0.040 | 16S | g__Paenibacillus |
| Otu001242 | >6 | shoots | 2.368 | 0.003 | 0.003 | 16S | g__Tissierella_Soehngenia |
| Otu001934 | >6 | shoots | 2.766 | 0.015 | 0.015 | 16S | g__Actinoplanes |
| Otu00007 | 6 | shoots | 2.734 | 0.017 | 0.017 | ITS2 | g__Acremonium |
| Otu00008 | >6 | shoots | 4.517 | 0.009 | 0.009 | ITS2 | g__Boeremia |
| Otu00090 | 6 | shoots | 3.435 | 0.025 | 0.025 | ITS2 | g__Nectriella |
| Otu00559 | 4 | shoots | 2.710 | 0.018 | 0.018 | ITS3 | f__Didymellaceae_unclassified |
| Otu01220 | <4 | shoots | 2.739 | 0.002 | 0.002 | ITS3 | g__Pseudoseptoria |
| Otu00294 | >6 | shoots | 3.251 | 0.008 | 0.008 | 18S | Panagrolaimus |
| Otu01388 | >6 | shoots | 2.312 | 0.002 | 0.002 | 18S | Panagrolaimus |
| Otu000112 | >6 | flowers | 3.019 | <0.001 | <0.001 | 16S | f__Planococcaceae_unclassified |
| Otu000140 | 4 | flowers | 2.855 | 0.002 | 0.002 | 16S | g__Frigoribacterium |
| Otu000276 | 4 | flowers | 2.684 | 0.010 | 0.010 | 16S | g__Nocardioides |
| Otu001033 | 4 | flowers | 2.284 | 0.001 | 0.001 | 16S | g__Chryseobacterium |
| Otu001273 | 4 | flowers | 2.237 | 0.032 | 0.032 | 16S | k__Bacteria_unclassified |
| Otu002511 | 6 | flowers | 2.041 | <0.001 | <0.001 | 16S | k__Bacteria_unclassified |
| Otu00007 | >6 | flowers | 2.733 | 0.006 | 0.006 | ITS2 | g__Acremonium |
| Otu00488 | 6 | flowers | 2.924 | 0.032 | 0.032 | ITS2 | k__Fungi_unclassified |
| Otu00610 | 5 | flowers | 3.457 | 0.039 | 0.039 | ITS2 | g__Cenococcum |
| Otu00558 | >6 | flowers | 3.549 | 0.011 | 0.011 | 18S | Eukaryota_unclassified |
| Otu00770 | 4 | flowers | 2.837 | 0.049 | 0.049 | 18S | Eukaryota_unclassified |
| Otu000074 | 4 | seeds | 2.651 | 0.024 | 0.024 | 16S | f__Sinobacteraceae_unclassified |
| Otu000122 | 4 | seeds | 2.185 | 0.015 | 0.015 | 16S | f__Gaiellaceae_unclassified |
| Otu000275 | >6 | seeds | 3.189 | 0.022 | 0.022 | 16S | f__Burkholderiaceae_unclassified |
| Otu000289 | <4 | seeds | 2.131 | 0.018 | 0.018 | 16S | g__Marmoricola |
| Otu000819 | <4 | seeds | 2.032 | <0.001 | <0.001 | 16S | o__Bacillales_unclassified |
| Otu001353 | 6 | seeds | 2.621 | 0.020 | 0.020 | 16S | g__Cupriavidus |
| Otu001638 | >6 | seeds | 3.030 | 0.036 | 0.036 | 16S | k__Bacteria_unclassified |
| Otu002021 | >6 | seeds | 2.835 | 0.003 | 0.003 | 16S | k__Bacteria_unclassified |
| Otu002029 | <4 | seeds | 2.063 | 0.037 | 0.037 | 16S | g__Mesorhizobium |
| Otu00058 | 5 | seeds | 2.276 | 0.010 | 0.010 | ITS2 | g__Colletotrichum |
| Otu01315 | <4 | seeds | 2.558 | 0.036 | 0.036 | ITS2 | k__Fungi_unclassified |
| Otu01406 | >6 | seeds | 2.772 | 0.008 | 0.008 | ITS2 | k__Fungi_unclassified |
| Otu01771 | <4 | seeds | 2.377 | 0.002 | 0.002 | ITS3 | k__Fungi_unclassified |
